# Altered M2 Cortex - Superior Colliculus visual perception circuit in Huntington’s Disease

**DOI:** 10.1101/2022.03.24.485610

**Authors:** Sara Conde-Berriozabal, Lia García-Gilabert, Esther García-García, Laia Sitjà-Roqueta, Javier López-Gil, Emma Muñoz-Moreno, Mehdi Boutagouga Boudjadja, Guadalupe Soria, Manuel J Rodríguez, Jordi Alberch, Mercè Masana

**Affiliations:** Department of Biomedical Sciences, Institute of Neurosciences, School of Medicine and Health Sciences, University of Barcelona, 08036 Barcelona, Spain; Institut d’Investigacions Biomèdiques August Pi i Sunyer (IDIBAPS), 08036 Barcelona, Spain; Centro de Investigación Biomédica en Red sobre Enfermedades Neurodegenerativas (CIBERNED), Madrid, Spain; Magnetic Resonance Imaging Core Facility, Institut d’Investigacions August Pi i Sunyer (IDIBAPS), Barcelona, Spain; Metofico ltd, London, United Kingdom; Laboratory of Surgical Neuroanatomy, Institute of Neurosciences, School of Medicine and Health Sciences, Universitat de Barcelona, Barcelona, Spain; Consorcio Centro de Investigación Biomédica en Red (CIBER) de Bioingeniería, Biomateriales y Nanomedicina (CIBER-BBN), Madrid, Spain

## Abstract

The premotor cortical area -M2 cortex in rodents- connection to the striatum is involved in movement and prominently affected in Huntington’s Disease (HD). M2 cortex also projects to the superior colliculus (SC), implicated in oculomotor functions (i.e. saccades) and visual perception. Here, we investigated the contribution of M2 cortex – SC circuit to HD physiopathology in male mice. Using fMRI, we observed that M2 cortex functional connectivity with the SC was the most prominently affected circuit in the symptomatic R6/1 HD mouse model. Structural alterations were also detected by tractography and viral tracing. HD mice showed decreased defensive behavioral responses towards an unexpected visual stimuli, such as a moving robo-beetle, and decreased locomotion upon unexpected flash of light. Additionally, optogenetic M2 cortex – SC stimulation promoted avoidance responses towards the robo-beetle in WT, but not HD mice. Though, DWI measurements in vivo and ex vivo electrophysiological responses to optogenetic and electrical stimuli in the SC suggested preserved brain structure. Finally, GCamp6f fluorescence recordings with fiber photometry indicated that aberrant M2 cortex engagement might be the underlying mechanism of visual perception alterations in HD. Collectively, our findings point to a key role of M2 cortex - SC circuit alterations in HD pathophysiology.

## Introduction

Huntington’s disease (HD) is a neurodegenerative disorder characterized by devastating motor symptoms, including chorea, and preceded by cognitive and psychiatric disturbances. HD pathology is caused by a CAG trinucleotide expansion (>36 repeats) in the huntingtin gene (HDCRG et al, 1993) and results in a progressive degeneration of medium-sized spiny neurons in the caudate and putamen (striatum). The cortex is the main input to the striatum and an early and progressive cortico-striatal disconnection has been extensively demonstrated by several studies in HD patients (Dogan et al., 2015; Unschuld et al., 2012) and animal models (Cepeda et al., 2007; Veldman and Yang, 2018). In HD patients, the most affected structural and functional striatal connexions arise from cortical premotor areas, analogous to the secondary motor (M2) cortex in rodents. Notably, these affectations appear many years before the onset of motor symptoms in HD carriers (Estevez-Fraga et al., 2021; Johnson et al., 2021; Shaffer et al., 2017; Unschuld et al., 2012) and are profoundly impaired in animal models (Creus-Muncunill et al., 2019; Fernández-García et al., 2020; Hintiryan et al., 2016). However, it is unclear whether dysregulated information flow from the M2 cortex in HD affects only cortico-striatal functions or could have an additional impact in the remaining long-distance output nuclei.

Cortico-striatal circuits are required for motor learning processes (Cao et al., 2015; Costa et al., 2004; Kupferschmidt et al., 2017) and M2 cortex projection to the striatum has been associated to the motor learning deficits that characterize HD mice (Fernández-García et al., 2020). Furthermore, M2 cortex is involved in a variety of functions including not only motor learning processes (Cao et al., 2015) but also decision-making (Sul et al., 2011) and perceptual behaviour (Duan et al., 2021), as widely reviewed (Barthas and Kwan, 2017; Yang and Kwan, 2021). In this regard, pyramidal tract neurons from M2 cortex project ipsilaterally to striatum and also send collateral axons to the thalamus, subthalamic nucleus, superior colliculus (SC), pons, and spinal cord (Economo et al., 2018; Gerfen et al., 2018; Zhang et al., 2016). Particularly, M2 cortex is one of the major inputs to the SC (Comoli et al., 2012; Savage et al., 2017).

The SC receives and integrates visual information to control reflex movements such as saccades (rapid eye movements towards/away visual stimulus) (Sparks, 1986), stimulus orienting (approach response) or defensive movements (avoidance or flight) (Dean et al., 1989; Sparks, 1986; Wurtz and Albano, 1980), and complex functions involving attention and decision making (Basso et al., 2021). Interestingly, deficits in visual perception and oculomotor functions, such as saccadic movement alterations, precede motor symptoms in movement disorders (Termsarasab et al., 2015). Indeed, alterations in saccadic movements are present in pre-manifest HD patients, serve as quantitative biomarkers and can aid in predicting onset and severity of disease (Antoniades et al., 2007; Patel et al., 2012; Winder and Roos, 2018). Yet, the involvement of the SC in HD pathology has not been directly explored.

In the present work we aimed to determine for the first time the contribution of M2 cortex - SC circuit to HD pathology, using the R6/1 HD mouse model. First, we mapped alterations in M2 cortex functional connectivity with the rest of the brain in wild-type and HD mice, by using in vivo resting state fMRI techniques. Then, we studied the structural connectivity of M2 cortex - SC pathway by both diffusion MRI analysis and viral tracing. To determine M2 cortex - SC functional alterations in HD mice, we took advantage of optogenetic tools, multi-electrode arrays (MEAs), fiber photometry and SC dependent behavioural responses to unexpected visual stimuli, such as the presence of a moving robo-beetle (Almada et al., 2018) and an unexpected flash of light (Liang et al., 2015). Collectively, our findings reveal that the structure and function of M2 cortex - SC circuit is deeply impaired in the R6/1 HD mouse model and highlight the importance of studying beyond basal ganglia circuits to fully understand the progression of brain circuit alterations in HD neuropathology.

## Results

### The M2 cortex is functionally disconnected with the SC in HD mice

To investigate M2 cortex connectivity alterations in HD, we used rs-fMRI and analyzed functional connectivity using seed-based analysis between the M2 cortex and 22 automatically identified brain regions covering the left brain hemisphere in ∼20-week-old wild-type and R6/1 mice (Figure 1; Supplementary files 1-2). M2 cortex shows the highest functional connectivity with Cg and SC in WT mice. In addition, functional connectivity between the M2 cortex and all brain regions analyzed was reduced in HD mice compared to WT. Two-way ANOVA reported a significant genotype effect (*F*_(1,22)_ = 11.7; *p*=0.002), brain region effect (*F*_(20,440)_ = 36.2, *p*<0.0001) and region/genotype interaction effect (*F*_(20,440)_ = 3.6, *p*<0.0001). The left M2 cortex of HD mice, compared to WT, showed significantly reduced functional connectivity with left periaqueductal gray (PAG; *p*=0.001), SC (*p*<0.0001), thalamus (*p*=0.05), hippocampus (*p*=0.01), RS (*p*=0.001), Cg (*p*=0.007), Som (*p*=0.03) and mPFC (*p*=0.02), as shown by Bonferroni post-hoc test. Thus, our results highlight that M2 cortex functional connectivity deficits in symptomatic HD mice involves several cortical, hippocampal and non-basal ganglia brain regions, with most prominent changes with the SC.

**Figure 1.**
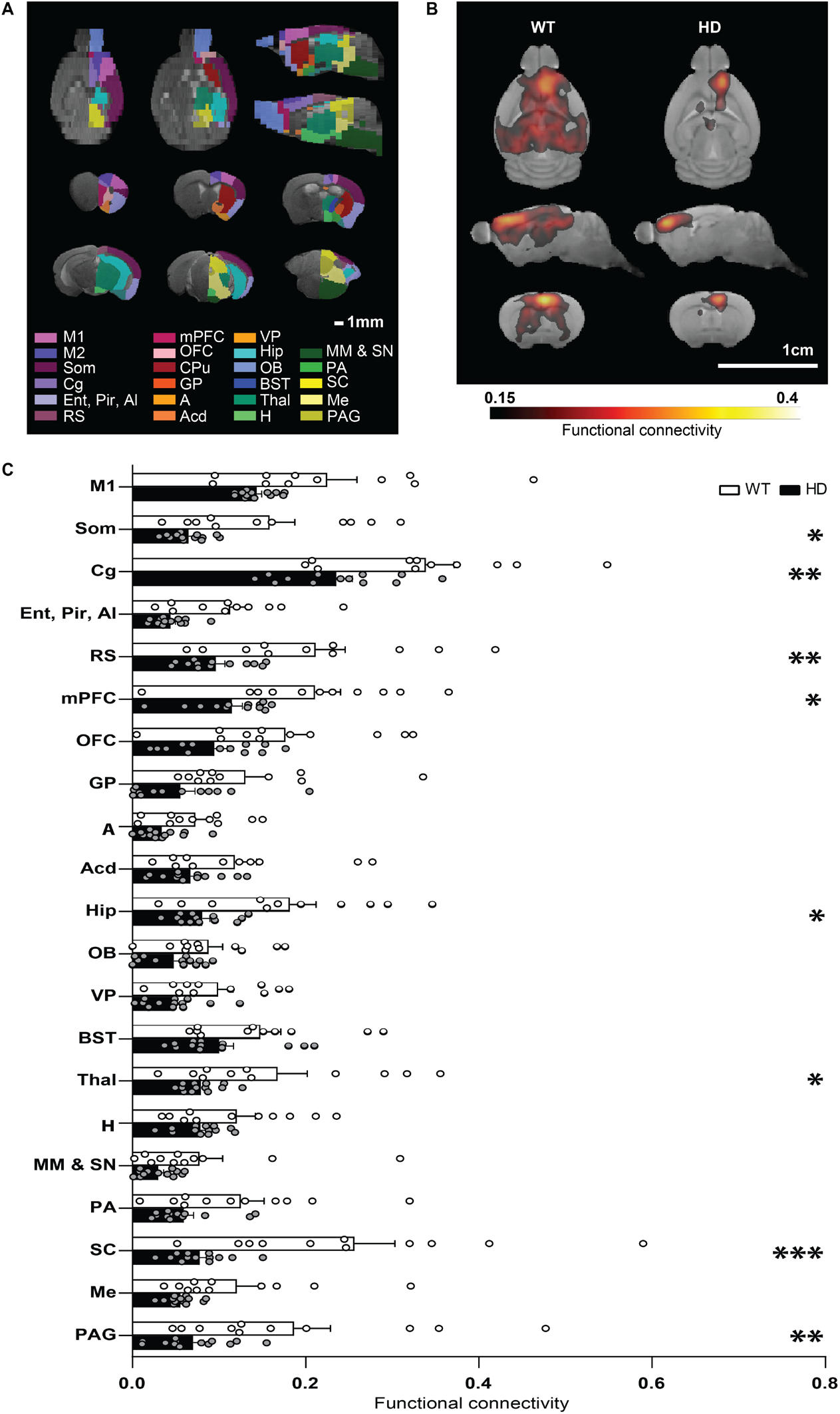
Functional connectivity analysis using rs-fMRI point the superior colliculus as the brain region most severely disconnected from M2 cortex in 20-week-old HD mice. **(A)** Anatomical regions for functional connectivity analysis based on atlas automatic parcellation. **(B)** Average functional connectivity maps of left M2 cortex in WT and the R6/1 mouse model of HD. Color maps represent the average correlation value when greater than 0.15. **(C)** Average functional connectivity of the left M2 cortex with all brain areas analyzed from the left hemisphere are represented. Average of the seed-based correlation map in each area represents functional connectivity between the M2 cortex and each region. Each point represents data from an individual mouse. Two-way ANOVA with genotype and brain region as factors was performed, followed by Bonferroni post-hoc comparisons test. Data are represented as mean ± SEM (WT n = 11 and HD n = 13 mice). \**p*<0.05, \*\**p*<0.01, \*\*\**p*<0.001 HD versus WT. Abbreviations: Primary motor (M1) cortex; Secondary motor (M2) cortex; Secondary somatosensory cortex (Som); Cingulate cortex (Cg); Entorhinal, Piriform and Agranular insular cortex (Ent, Pir, AI); Retrosplenial cortex (RS); Medial prefrontal cortex (mPFC); Orbitofrontal cortex (OFC); Caudate putamen (CPu); Globus pallidus (GP); Amygdala (A); Nucleus accumbens (Acd); Hippocampus (Hip); Olfactory bulb (OB); Ventral pallidum (VP); Bed nucleus of the stria terminalis (BST); Thalamus (Thal); Hypothalamus (H); Medial mammillary nucleus and Substantia nigra (MM & SN); Preoptic area (PA); Superior colliculus (SC); Mesencephalic nucleus (Me); Periaqueductal gray (PAG). **Supplementary file 1.** Average seed-based BOLD correlation maps from WT mouse M2 cortex related to Figure 1b, visualized with ITK-SNAP software (Yushkevich et al., 2006). **Supplementary file 2.** Average seed-based BOLD correlation maps from R6/1 HD mouse M2 cortex related to Figure 1b, visualized with ITK-SNAP software (Yushkevich et al., 2006).

### Structural alterations in M2 cortex - SC circuit of HD mice

To evaluate if functional connectivity deficits observed in M2 cortex - SC circuit were associated to structural alterations, we estimated the M2 cortex - SC tracts from DWI and measured FA as well as MD, AD and RD diffusivity (Figure 2). Our data indicates that structural alterations are also present in the M2 cortex - SC circuit. We reported significant decreased FA values (Figure 2B) when comparing HD with WT mice, while MD, AD and RD remains similar between genotypes (Figure 2C-E). In addition, we analysed DTI-metrics in the SC to exclude possible regional microstructural alterations (Figure 2-figure supplement 1). All the evaluated diffusion metrics from the SC were similar between WT and HD mice, suggesting that alterations in the M2 cortex - SC are related to the circuit activity, not the structure per se.

**Figure 2.**
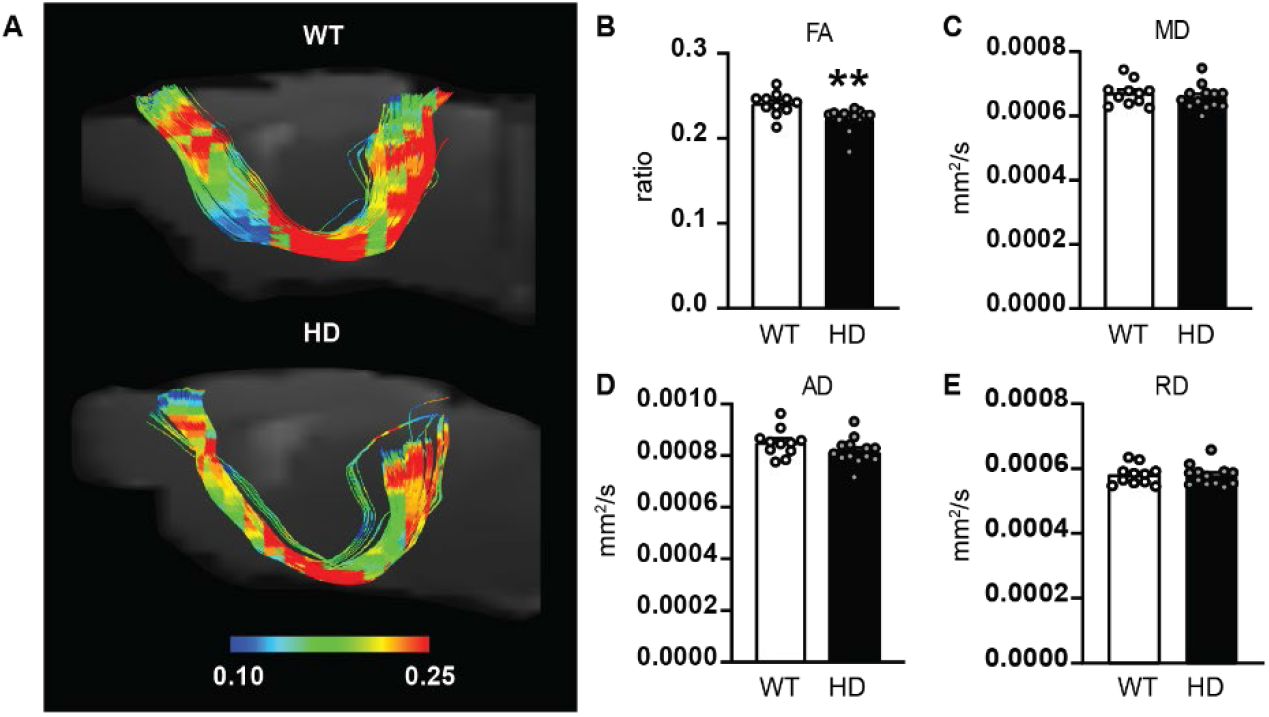
Diffusion metrics indicate structural alterations in M2 cortex – SC tracts of HD mice. **(A)** Representative tracts connecting the M2 cortex area with the SC in WT and the R6/1 mouse model of HD. M2 cortex and SC areas were automatically delimited according to atlas. **(B-E)** Diffusion metrics analysis of tracts between M2 cortex and SC includes **(B)** FA, **(C)** MD, **(D)** AD and **(E)** RD. Each point represents data from an individual mouse. Unpaired Student’s t-test was performed. Data are represented as mean ± SEM (WT n = 11 and HD n = 13 mice). \**p*<0.05, \*\**p*<0.01.

**Figure 2-figure supplement 1.**
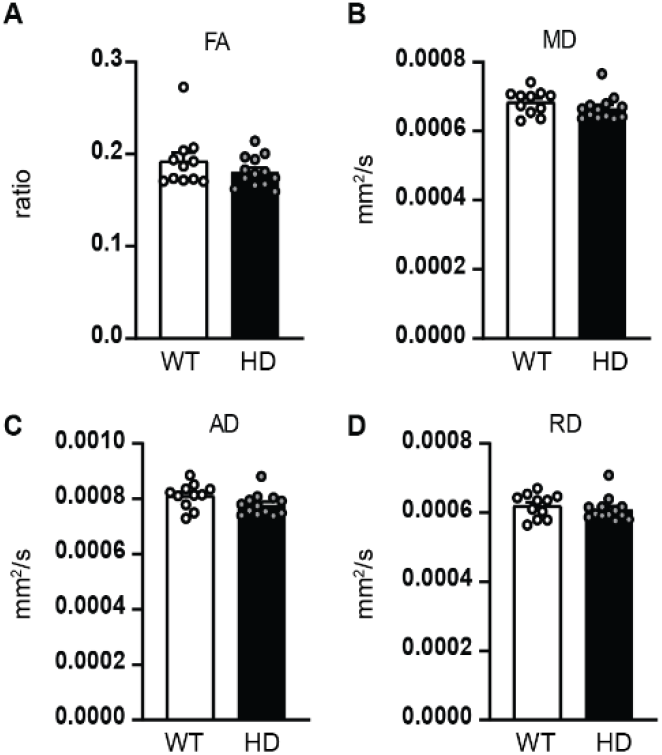
Regional diffusion metrics indicate preserved microstructure in the SC of HD mice. (A) FA, (B) MD, (C) AD and (D) RD were measured in the SC of 20-week-old WT and R6/1 mice. Each point represents data from an individual mouse. Unpaired Student’s t-test was performed. Data are represented as mean ± SEM (WT n = 11 and HD n = 13 mice).

Then, we took advantage of tracing techniques to further evaluate microstructural alterations in M2 cortex - SC circuit. Injections into the M2 cortex of an AAV-CamKII-YFP construct produced dense YFP expression in pyramidal neurons, whose terminals reach the deep layers of the lateral part of the of the SC (dlSC) (Figure 3), consistent with previously published work (Comoli et al., 2012; Duan et al., 2021; Oh et al., 2014; Savage et al., 2017). We quantified fluorescent intensity in the dlSC and observed significant reduction in YFP mean intensity values of HD mice, when compared to WT (Figure 3C). Our data confirms structural alterations in the M2 cortex - SC circuit in HD, with reduced axonal density arising from M2 cortex reaching the SC.

**Figure 3.**
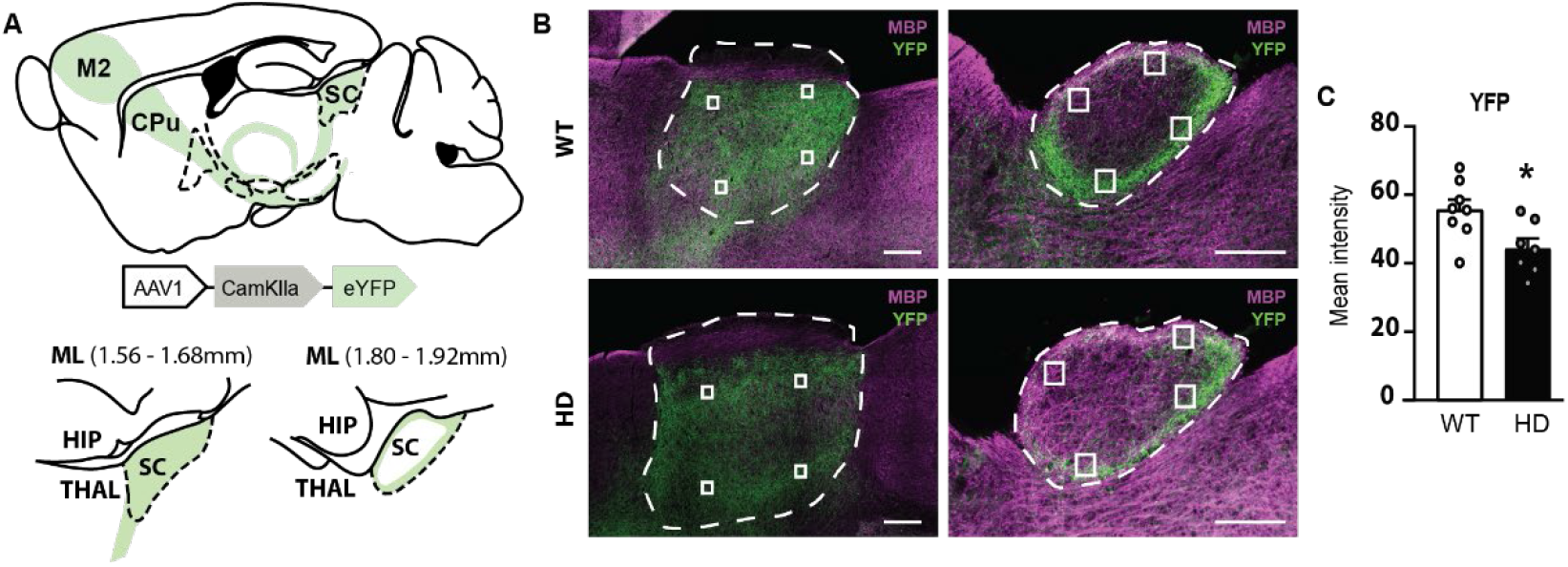
Injection of AAV-CamKII-YFP in M2 cortex induces reduced fluorescent axonal signal in the lateral superior colliculus of symptomatic HD mice. **(A)** Schematic representations showing AAV-CamKII-YFP construct, the location of injection at M2 cortex and its labelled projections and M2 cortical projections of two medio-lateral (ML) coordinates of the dlSC. **(B)** Representative fluorescent images showing M2 cortical projections expressing YFP (green) and myelin sheaths stained with Myelin Basic Protein (MBP) (pink) in the dlSC. Scale bar: 250 µm. Dotted lines: SC area. Squares: Examples of manually selected ROIs for quantification. **(C)** Mean fluorescence intensity of M2 cortex YFP labelled axons in SC. Data are represented as mean ± SEM (4 ROI / slice; WT n = 8 and HD n = 7 slices). \**p*<0.05, unpaired Student’s t-test.

### Impaired behavioural function of the SC in symptomatic HD mice

The SC integrates visual information to generate stimulus orienting or defensive responses. To examine if SC dependent behaviours were affected in HD mice, we selected two tasks known to involve SC function, the BMT (Almada et al., 2018) and the Looming task (Liang et al., 2015).

In the BMT (Figure 4 and Video 1), the behavioral responses to a randomly moving robo-beetle are evaluated. During the habituation phase of the test, mice were allowed to explore the arena for 5 min and locomotion and spontaneous exploration such as rearing time was significantly decreased in HD mice compared to WT (Figure 4B-C), as previously described (Pépin et al., 2016). During the testing phase, when mice were confronted with the robo-beetle, HD mice showed a significant decrease in total number of contacts with the robo-beetle (Figure 4D). Then, we evaluated the type of response towards the robo-beetle contact, and we observed that the major response in WT mice was to escape from the robo-beetle, compared to the other evaluated responses such as tolerance, approach or avoidance. Interestingly, HD mice show little or non-escape response, as demonstrated by the strongly reduced total percentage of escape response (Figure 4E). We further evaluated the percentage of responses when excluding escape behavior, as previously done in (Almada et al., 2018) and found that number of total encounters between HD mice and the robo-beetle was significantly higher compared with WT when obviating escape responses (Figure 4F). Of those, HD mice had significantly increased both tolerance (Figure 4G) and approach responses (Figure 4H). Finally, although avoidance behaviour is the preferred response to the robo-beetle in both genotypes (when excluding escape responses), HD mice showed significantly reduced avoidance responses compared to WT mice (Figure 4I). Altogether, our data highlights that the choice of response to unexpected threatening beetle differs in HD compared WT mice, suggesting alterations in visual perception processes involving the SC.

**Figure 4.**
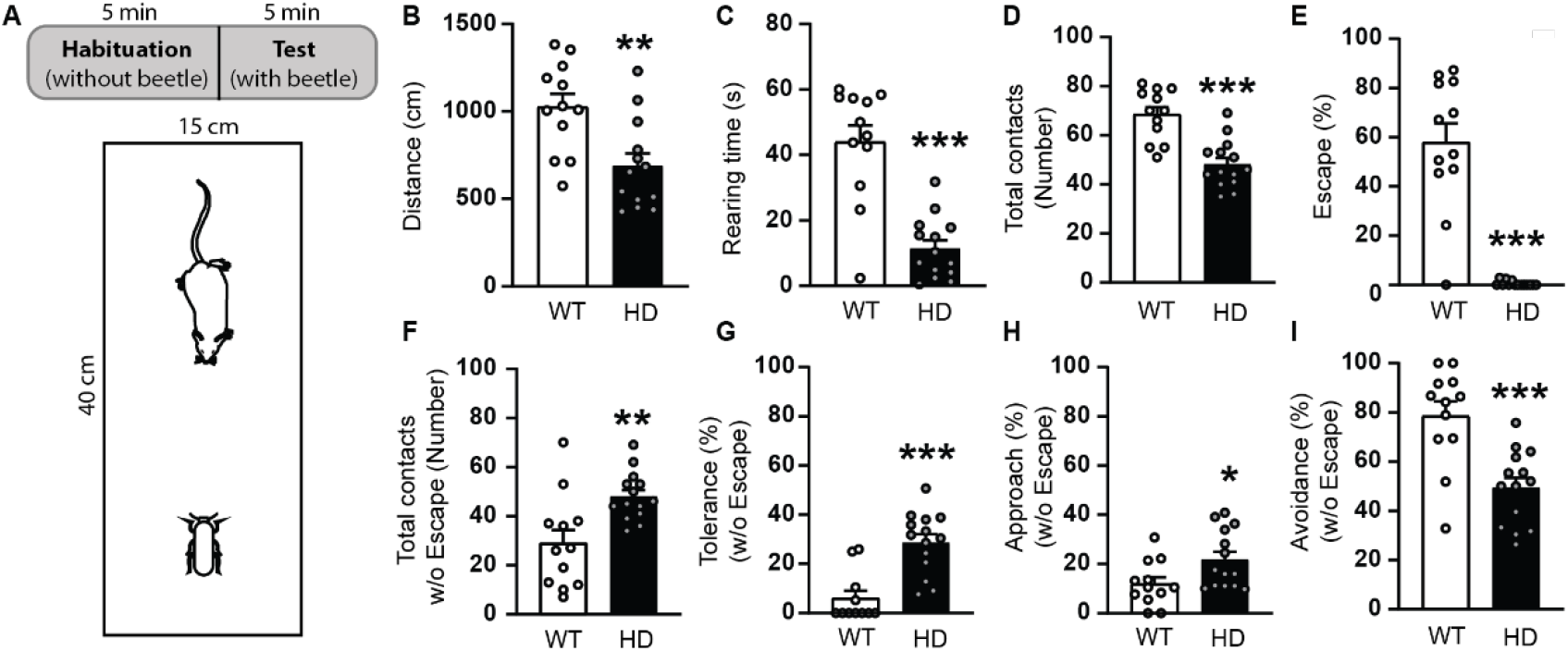
Behavioral responses towards a randomly moving robo-beetle differ between WT and HD mice at symptomatic stages. **(A)** Schematic representation of the BMT protocol, including 5 min habituation and 5 min test phases in a rectangular arena. **(B-C)** During the habituation phase **(B)** distance travelled and **(C)** rearing time were measured. **(D-I)** During the test phase, **(D)** total contacts with the robo-bettle and percentage of induced response are represented. Percentages of **(E)** escape responses, **(F)** total contacts excluding escape and percentage of **(G)** tolerance, **(H)** approach and **(I)** avoidance responses when excluding the escape response were analyzed. Each point represents data from an individual mouse. Unpaired Student’s t-test was performed. Data are represented as mean ± SEM (WT n = 12 and HD n = 14 mice). \**p*<0.05, \*\**p*<0.01. **Video 1. Representative videos of a WT and an HD mouse behavior in the presence of a robo-beetle during the BMT.**

Next, we assessed light-triggered behaviours in response to unexpected stimuli from the upper-visual field using the Looming task (Liang et al., 2015). We compared the response to a sudden flash of white light (∼1 s duration) in the center of a dark corridor between genotypes (Figure 5). We observed that a sudden light induced a short freezing in WT mice (data not shown), although we did not observe reduction in time to travel through the corridor after the light in WT mice as previously described (Liang et al., 2015), we found that time was increased in HD mice. Two-way ANOVA reported a significant genotype effect (*F*_(1,24)_ = 11.0, *p*=0.003), time 1/2 effect (*F*_(1,24)_ = 20.8, *p*=0.0001) and interaction effect (*F*_(1,24)_ = 10.9, *p*=0.003). Bonferroni post-hoc analysis showed that locomotion was not affected in WT mice by the sudden light (Figure 5B), whereas it was significantly increased in HD mice and consistent among trials (Figures 5C-D). On the one hand, time to reach the center of the corridor was affected by the trial number, as shown by two-way ANOVA with a significant trial effect (*F*_(4,85)_ = 6.8, *p*<0.0001), but neither genotype effect (*F*_(1,24)_ = 4.3, *p*=0.05) nor interaction (*F*_(4,85)_ = 6.8, *p*=0.09) (Figure 5C). On the other hand, time from the center of the corridor to the end was increased in all trials in HD mice, as shown by two-way ANOVA significant genotype effect (*F*_(1,24)_ = 15.8, *p*=0.0006) and interaction (*F*_(4,85)_ = 3.8, *p*=0.007), but no trial effect (*F*_(4,85)_ = 1.1, *p*=0.3) (Figure 5D). Therefore, our data further confirm that WT and HD mice generate divergent responses in response to unexpected visual stimuli, indicating that SC function is affected in HD.

**Figure 5.**
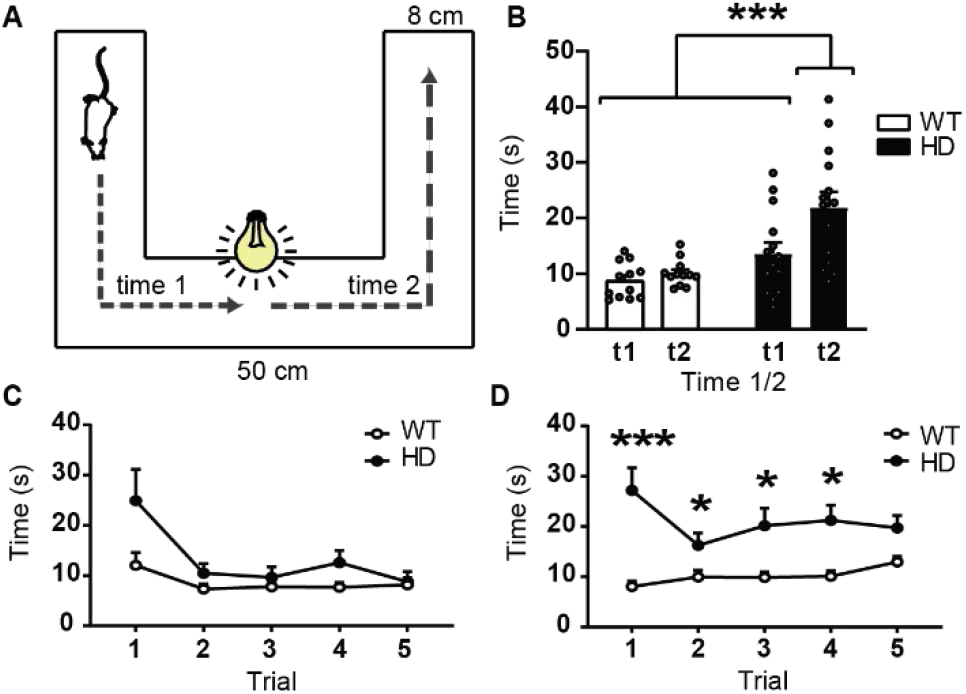
The behavioural response to unexpected flash of light is altered in symptomatic HD mice. **(A)** Schematic representation of the Looming task protocol. Mouse will travel through a corridor and a sudden flash of light is opened when reaches the centre. **(B)** Plot of average time to travel until the centre (Time 1) and from the centre to the end of the corridor (Time 2). Each point represents the average of the 5 trials from an individual mouse. **(C-D)** Graphs showing the measures of time 1 **(C)** and time 2 **(D)** over the 5 trials. Two-way ANOVA genotype and time or trials as factors was performed and followed by Bonferroni post-hoc test comparisons. Data are represented as mean ± SEM (WT n = 12 and HD n = 14 mice). \**p*<0.05, \*\**p*<0.01, ***p<0.001.

### M2 cortex inputs to the SC promotes avoidance behavior in WT but not HD mice

In order to dissect the involvement of M2 cortex inputs to the behavioral responses associated to the SC, we induced the activity of the M2 cortical afferents to the SC using optogenetics. Because neurons from the M2 cortex project to the dlSC and are associated to lower visual fields (Comoli et al., 2012), we evaluated the responses of a lower visual stimulus, such as the robo-beetle, during optogenetic stimulation in an extended version of the BMT (Figure 6). We expressed ChR2 specifically in M2 cortical pyramidal neurons by injecting an AAV-CamKII-ChR2-YFP construct and modified the BMT protocol by prolonging to 10 min the testing phase. Selective stimulation of the M2 cortical afferents in the SC was performed during the last 5 min of the test. Similar to prior results, we found a significant decrease in both locomotion (Figure 6-figure supplement1A) and time spent exploring the arena (Figure 6-figure supplement1B) during the habituation phase in HD mice compared to WT. During the subsequent testing phase, we quantified the behavioural responses towards the robo-beetle, before and during activation of the M2 cortex - SC circuit. In line with previous results, the total number of contacts was reduced in HD mice (Figure 6D), reported by two-way ANOVA genotype effect (*F*_(1,10)_ = 6.4, *p*=0.03). Also, the escape response was the preferential response to the robo-beetle for WT mice, while almost none of HD mice performed this response two-way ANOVA genotype effect (*F*_(1,9)_ = 46.1, *p*<0.0001; Figure 6E). We found no stimulation nor interaction effects in these two parameters.

**Figure 6.**
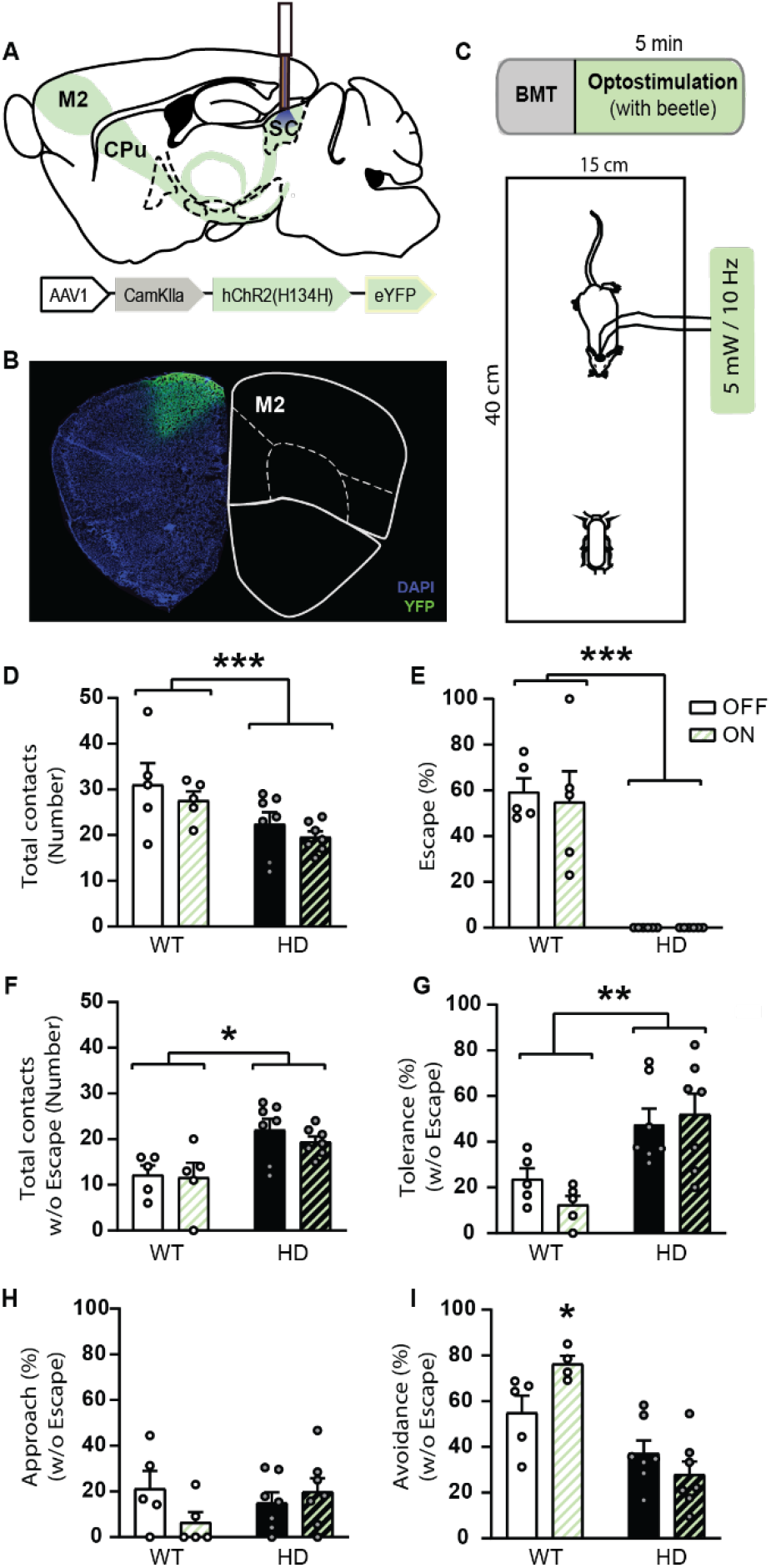
Optogenetic stimulation of M2 cortex terminals in the SC promotes avoidance behaviour in WT mice in the Beetle Mania Task, but not in HD mice. **(A)** Schematic representation of fiber-optic cannula implant in the dlSC, AAV-CamKII-ChR2-YFP construct and injection location at M2 cortex. **(B)** Representative fluorescent image showing DAPI (blue) and M2 cortical neurons expressing YFP (green). **(C)** Modified BMT protocol, including 5 min habituation, 5 min test plus 5 min test with optogenetic stimulation. **(D-I)** Total number of contacts and percentage of mice responses were analyzed during the first 5 min of test (Laser OFF) and then during the 5 min of test with optostimulation (Laser ON). **(D)** Total contacts with the robo-beetle. **(E)** Percentages of escape responses. **(F)** Total contacts excluding escape. **(G)** Percentage of tolerance, **(H)** approach and **(I)** avoidance responses, excluding the escape response. Each point represents data from an individual mouse. Two-way ANOVA with genotype and stimulation effects was performed and followed by Bonferroni post-hoc test comparisons when appropriate. Data are represented as mean ± SEM (WT n = 4/5 and HD n = 6/7 mice). \**p*<0.05, \*\**p*<0.01, \*\*\**p*<0.001 versus WT.

**Figure 6-figure supplement1.**
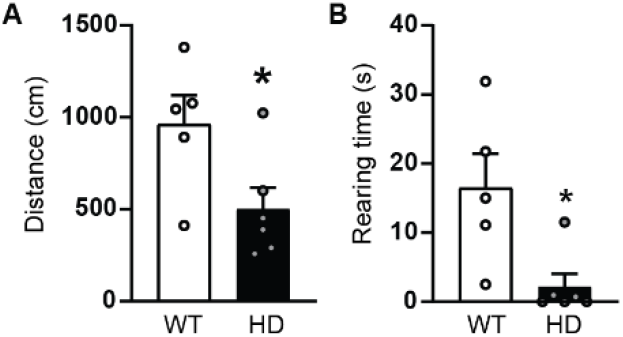
Locomotion and exploratory behaviour are reduced in the HD mice model of HD. Distance travelled (A) and rearing time (B) were measured during the 5 min of habituation of the BMT in 20-week-old R6/1 mouse model of HD. Each point represents data from an individual mouse. Unpaired Student’s t-test was performed. Data are represented as mean ± SEM (WT n = 5 and HD n = 7 mice). \**p*<0.05.

When excluding the escape responses, the total number of contacts was again increased in HD mice two-way ANOVA genotype effect (*F*_(1,10)_ = 10.0, *p*=0.01; Figure 6E), although neither effects of the optogenetic stimulation nor interaction effect were found. Tolerance responses were increased in HD mice, two-way ANOVA genotype effect (*F*_(1,10)_ = 11.9, *p*=0.006; Figure 6G) but we found no stimulation effect nor genotype/stimulation interaction. Nevertheless, we observed a tendency to reduction in WT-stimulated mice. Approach behaviour was similar between genotypes and was also not modulated by optogenetic stimulation, although we observed genotype/stimulation interaction two-way ANOVA interaction (*F*_(1,10)_ = 5.7, *p*=0.04; Figure 6H). Finally, avoidance responses (Figure 6I) were reduced in HD mice compared to WT mice, and optogenetic stimulation of M2 terminals was able to induce an increase of avoidance responses in WT mice, as demonstrated by two-way ANOVA of genotype effect (*F*_(1,10)_ = 23.1, *p*=0.0007) and genotype/stimulation interaction (*F*_(1,9)_ = 10.2, *p*=0.01). Bonferroni post-hoc analysis reported a significant increase in avoidance responses in WT mice in response to the optogenetic stimulation of M2 terminals in the SC, which did not occur in HD mice.

Altogether, our data demonstrates a direct involvement of M2 cortex - SC circuit in the selection of responses during the presence of a visual stimulus, showing enhanced choice of avoidance responses in WT mice. Additionally, our results further demonstrate that M2 cortex - SC circuit is deeply impaired in HD, as optogenetic stimulation failed to modulate responses in those mice.

### Stimulation of M2 cortical axons in the SC induces electrophysiological responses in WT and HD mice

To elucidate whether acute optical stimulation was efficiently activating the M2 cortex - SC circuit in both WT and HD mice, we performed MEA recordings in the SC in WT and HD mice previously injected in the M2 cortex with the AAV-CamKII-ChR2-YFP construct (Figure 7). A normalized input-output assay was created by recording fPSC in the dlSC induced by increasing light intensities (Figure 7C). Our results revealed that optical stimulation of M2 cortical axons in the dlSC evoked fPSC in both WT and HD mice. Two-way ANOVA showed a significant light intensify effect (*F*_(7,84)_ = 34.5, *p*<0.0001) although neither genotype effect (*F*_(1,12)_ = 3.0, *p*=0.11) nor genotype/light intensity interaction (*F*_(7,84)_ = 1.9, *p*=0.08) were found. Bonferroni post-hoc analysis showed that the amplitude of the evoked fPSC in the dlSC was significantly greater in HD compared with its respective WT at the highest light intensity (79 mW) (*p*=0.002). These results validate our optogenetic approach and suggest that alterations in SC dependent behavioural functions are not due to deficient dlSC responses to M2 cortex stimulation. Additionally, we performed an input-output assay using increasing electrical stimulation in the dlSC (Figure 7D). Responses in the dlSC increased along the intensity of the electrical stimulation as expected and demonstrated by two-way ANOVA significant intensify effect (*F*_(10,120)_ = 18.9, *p*<0.0001) but were not affected by the genotype.

**Figure 7.**
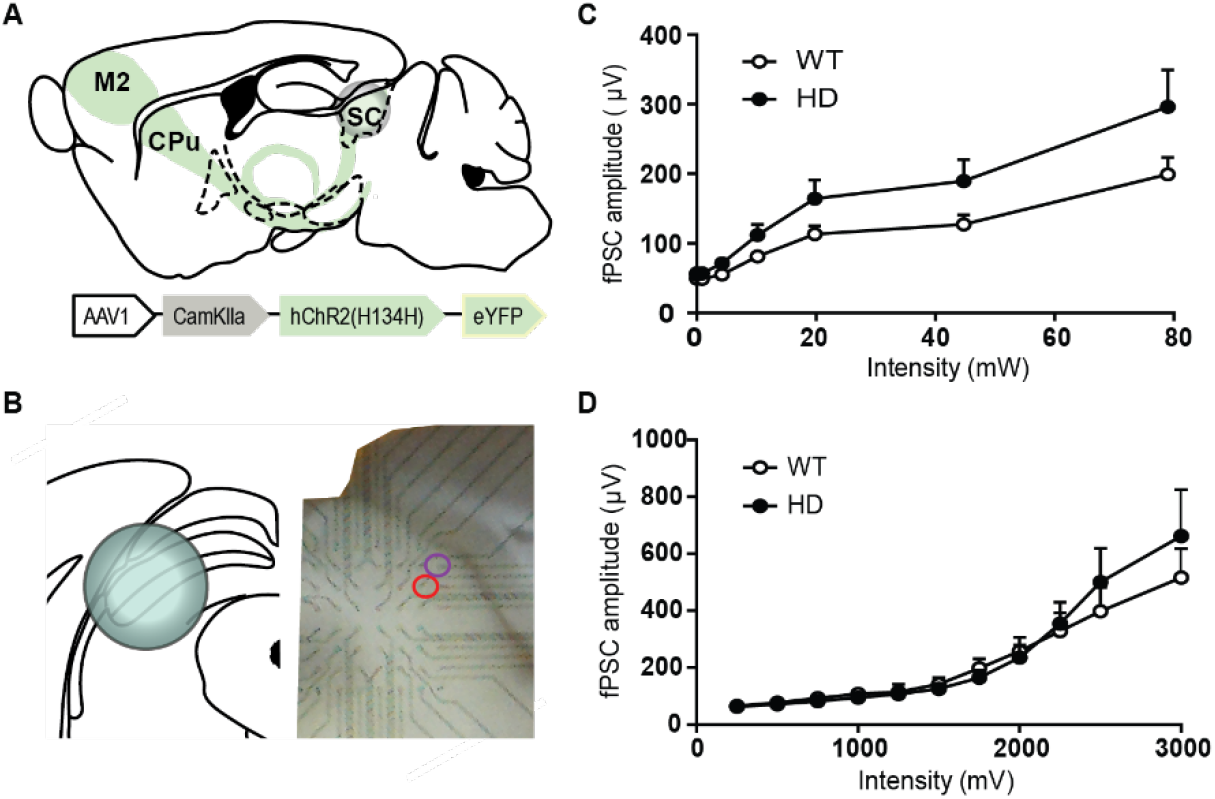
Optogenetic stimulation of M2 cortex afferents in the superior colliculus induces neuronal responses in both WT and HD mice, measured by multi-electrode array in brain coronal slices. **(A)** Schematic representation of AAV-ChR2 construct and injection location at M2 cortex and **(B)** MEA recordings in coronal slices. A fiber-optic cannula was placed on top of the dlSC and used for light stimulation. One electrode (purple circle) in the dlSC was used for electrical stimulation. Examples of a stimulating and recording (red circle) electrodes are shown in the representative image. **(C)** Amplitude of light intensity-induced SC fPSC in WT and HD mice. **(D)** Amplitude of the electrically-induced fPSC in the SC of WT and HD mice. Data are represented as mean ± SEM (WT n = 7 and HD n = 8 mice).

### Visual stimuli fails to engage neuronal activity in M2 cortex of HD mice

Our previous data indicate that SC neurons can respond similarly to M2 cortical inputs in both WT and HD mice. We therefore hypothesize that the function of M2 cortex neurons might be responsible of the SC-related alterations in HD mice. To prove that, we assessed whether M2 cortex activity correlated with the presence of an unexpected lower-visual stimuli (Figure 8). We expressed a GCaMP6f calcium sensor in M2 cortical neurons by injecting an AAV-GCaMP6g-WPRE-SV40 construct (Figure 8A-B) and recorded fluorescence during the 10 min of the BMT (Figure 8C). Fluorescence signal was stable and similar between genotypes during the habituation phase, and markedly increased straight after introducing the robo-beetle in WT mice, but not in HD mice (Figure 8D). To better quantify these results, we averaged fluorescent data, to obtain one min windows, and evaluated genotypes differences over time. Two-way ANOVA analysis reported a significant time effect (*F*_(5,45)_ = 4.7, *p*=0.002), genotype effect (*F*_(1,9)_ = 7.4, *p*=0.02) and time/genotype interaction (*F*_(5,19)_ = 3.1, *p*=0.02) (Figure 8E). Bonferroni post-hoc analysis showed significant differences between groups during the first three minutes of the testing phase. Thus, our data confirms that M2 cortex becomes active in WT mice in the presence of an unexpected lower-visual stimuli, but this neuronal engagement is lacking in HD mice.

**Figure 8.**
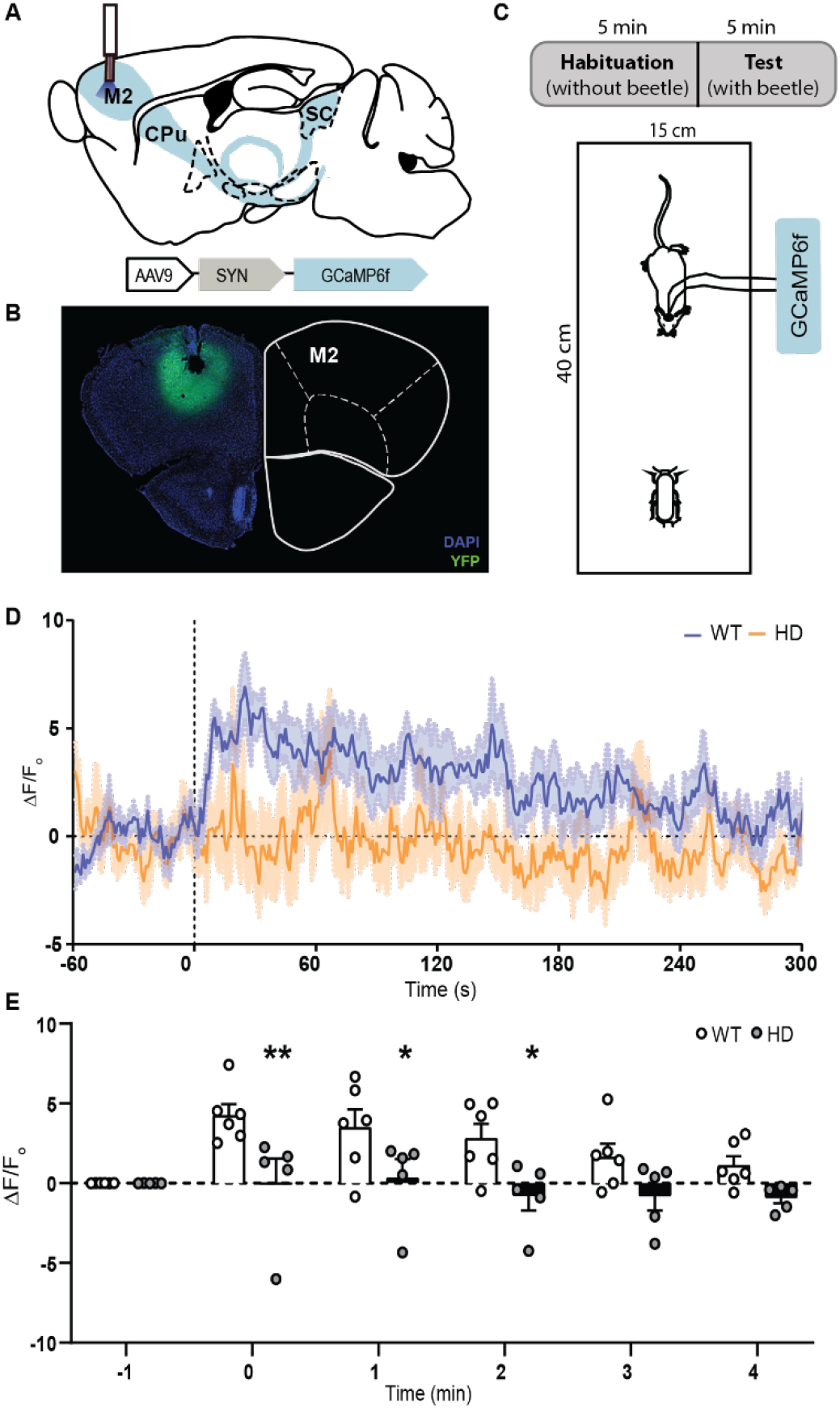
The presence of visual stimuli activates M2 cortex neurons in WT but not HD mice, shown by fluorescent calcium recordings using fiber photometry in vivo. **(A)** Schematic representation of fiber-optic cannula implant and representative image of the injection of the AAV-GCaMP6g-WPRE-SV40 construct at M2 cortex. **(B)** Representative fluorescent image showing DAPI (blue), YFP expression (green) and fiber-optic cannula implant in the M2 cortex. **(C)** BMT protocol, including 5 min habituation and 5 min test. **(D)** Mean photometric recordings of M2 cortex fluorescent calcium signal 1 min before and 5 min after the introduction of a moving robo-beetle to the arena in WT and symptomatic HD mice. Increases in fluorescent signal is normalized by the respective baseline for each mouse. **(E)** Average of normalized fluorescence signal is computed and represented for each minute. Each point represents data from an individual mouse. Two-way ANOVA analysis with genotype and time as factors was performed and followed by Bonferroni post-hoc test comparisons. Data are represented as mean ± SEM (WT n = 6 and HD n = 5 mice). \**p*<0.05, \*\**p*<0.01 versus WT.

## Discussion

Cortical premotor areas are prominently affected in HD many years before the onset of motor symptoms. These alterations have been widely correlated to cortico-striatal disconnection and associated to motor deficits, which are severely manifested in HD. Here, we thoroughly mapped M2 cortex circuit alterations in HD male mice and described severe structural and functional defects in the M2 cortex projection to the SC. Moreover, our data reveal profound alterations in SC dependent behavioural functions in HD mice, in agreement with the visual perception and oculomotor dysfunctions described in HD carriers (Antoniades et al., 2007; Patel et al., 2012; Winder and Roos, 2018). Finally, our results suggest that the underlying mechanism of these alterations in HD might be a lack of M2 cortex engagement upon visual stimuli.

M2 cortex pyramidal neurons send projections to many cortical and subcortical brain nuclei, including the SC, as previously described (Economo et al., 2018; Gerfen et al., 2018; Lin et al., 2018; Zhang et al., 2016). Notably, fMRI shows sever deficits in functional connectivity between the M2 cortex and the SC in HD, which is accompanied by structural circuit alterations, seen by a decrease in DWI metrics and viral tracing. Thus, we demonstrate that long-range pyramidal tract neurons from M2 cortex are severely affected, which contrasts with the idea that intratelencephalic neurons are more affected in HD (Gatto et al., 2015). Consistently, specific optogenetic activation of M2 cortical axons in the dlSC modulates mice responses to visual stimuli in the BMT in control, but not HD mice, while SC structure seems preserved in HD conditions, as shown by DWI measurements in the SC and MEA recordings. Overall, our data suggest that alterations in M2 cortex - SC functions are related to M2 cortex inputs rather than deficits in the SC structure. Whether additional inputs to the SC are also affected remains unknown.

In this regard, M2 cortex display the highest functional connectivity with the Cg, which in turn is the main cortical afferent to the medial portion of the SC (mSC) (Savage et al., 2017). Information flows from M2 cortex and Cg to the SC in functionally segregated loops and then goes back to basal ganglia circuitry through distinct thalamic targets to modulate action selection (McHaffie et al., 2005). In detail, M2 cortex - dlSC pathway function is related to approach and appetitive stimuli normally found in the lower visual field (Dean et al., 1986; Sahibzada et al., 1986), while Cg - mSC function is more related to escape behaviours and movements in the upper visual field (e.g., predators) (Comoli et al., 2012; Savage et al., 2017). Consequently, the presence of alterations in all the diverse behavioural responses in HD mice suggest that inputs to both the medial and lateral parts of the SC might be affected. Accordingly, Cg is known to be altered in HD patients at early stages (Hobbs et al., 2011), and cell loss in the motor and Cg correlates with symptomatology in HD (Thu et al., 2010). Therefore, we hypothesise that alterations in Cg - SC might also have a role in HD.

Additionally, M2 cortex - PAG circuit is also functionally altered in HD, as shown by the fMRI data. The PAG is one of the major outputs of the SC and is responsible of defining the threshold to compute escape responses (Branco and Redgrave, 2020; Evans et al., 2018), which is almost absent in HD conditions. Interestingly, increased GABAergic tone from the ventral lateral geniculate nucleus (Fratzl et al., 2021) or the substantia nigra reticulata (Almada et al., 2018) reduces escape responses in the BMT, suggesting that increased GABAergic activity in the SC could lead to the altered behavioural responses in HD mice. Further, SC - PAG circuit is shaped by multimodal sensory information and learning experiences, which are constantly updated throughout the life of the animal (Evans et al., 2019). However, whether M2 cortex could modulate PAG function through its few direct projections (Savage et al., 2017) or indirectly through the modulation of SC activity requires further investigation.

Our data highlights that M2 cortex is strongly engaged during the presence of a unexpected visual stimuli such as the robo-bettle, in accordance with its involvement in perceptual behavior (Duan et al., 2021). Furthermore, our results in HD mice suggest that the failure to engage M2 cortex activity may be the underlying mechanism to explain visual perception alterations. In this line, M2 cortex activation has been involved in many naturalistic behaviours such as rearing, grasping, eating, grooming etc, as seen by calcium imaging experiments using miniature microscopes (Tombaz et al., 2020) and to motor learning processes using two-photon imaging of Arc-GFP expression (Cao et al., 2015), and which behaviours are known to be altered in HD mice (Fernández-García et al., 2020; Pépin et al., 2016). A recent publication shows that inhibiting M2 terminals in the SC (using DREADDS) selectively impairs decision maintenance in a memory-dependent perceptual decision task, indicating an additional role of this circuit in motor planning (Duan et al., 2021). We therefore suggest that M2 cortex lack of engagement could be a common underlying mechanism in diverse HD symptoms.

Finally, symptomatic HD mice display impaired SC-related behaviours, which is consistent with the abnormal control of saccadic eye movements present in HD patients (Patel et al., 2012; Srivastava et al., 2018). Indeed, oculomotor functions and locomotion share many underlying neuronal circuits (Srivastava et al., 2018) and, in several movement disorders, oculomotor deficits precede motor symptoms (Termsarasab et al., 2015). Whereas direct alterations of SC function in HD patients have not been explored, studies in Parkinson Disease (PD) show that responses to luminescence (measured by fMRI) in the SC are abnormal in de novo patients compared to controls (Moro et al., 2020). These alterations in the SC are linked with deficits in the coordination of action and perception (Pretegiani et al., 2019). Notably, a recent study with PD patients show that repetitive transcranial magnetic stimulation in the motor cortex improved anti-saccade success rate and postural instability gait difficulty (Okada et al., 2021). In this line, optogenetic stimulation of the M2 cortex reverts motor dysfunction in a mouse model of PD (Magno et al., 2019). How visual perception in the SC modulates basal ganglia function is only starting to be understood. Thus, further understanding on the complex circuitry between cortical, visual and motors domains could help to design new treatment opportunities based on circuit restoration from early stages. While oculomotor dysfunction is present in both male and female HD patients (Patel et al., 2012), recent data indicate gender differences regarding HD progression (Zielonka and Stawinska-Witoszynska, 2020). Therefore, future studies including both genders are relevant, especially at early stages of disease.

In conclusion, our data provides compelling evidence of a key role of M2 cortex - SC circuit in HD pathophysiology, which might be also relevant for other movement disorders such as PD. Also, we highlight the lack of M2 cortex engagement as the underlying mechanism in HD visual perception symptoms. Thus, unravelling the myriad of circuit alterations and functions from M2 cortex other than cortico-striatal provides valuable insights to understand HD symptoms. Indeed, targeting visual perception and oculomotor circuits may provide novel therapeutic opportunities aiming to delay the onset and severity of symptoms in movement disorders.

## Materials and methods

### Key Resources Table

| Key Resources Table |  |  |  |  |
| --- | --- | --- | --- | --- |
| Reagent type (species) or resource | Designation | Source or reference | Identifiers | Additional information |
| Genetic reagent (Mus musculus) | B6CBA-Tg(HDexon1)61 Gpb/1J (R6/1) | Jackson laboratory | RRID:IMSR_JA X:002809 | R6/1 HD model |
| Recombinant DNA reagent | AAV1-CamKIIa-hChR2(H134H)-eYFP-WPRE.hGH | University of Pennsylvania-Penn Vector Core | Catalog number: AV-1-26969P | titres: $8.97 \times 10^{12}$ genomic particles/mL (we used 1:10 dilution) |
| Recombinant DNA reagent | AAV1-CamKIIa(1,3)-eYFP.WPRE.hGH; | University of Pennsylvania-Penn Vector Core | Catalog number: AV-1-PV2975 | titres: $1.18 \times 10^{13}$ genomic particles/mL (we used 1:10 dilution) |
| Recombinant DNA reagent | AAV9-Syn-GCaMP6f-WPRE-SV40 | University of Pennsylvania-Penn Vector Core | Catalog number: 100837-AAV9 | Titres: $1 \times 10^{13}$ genomic particles/mL |
| Antibodies | Rabbit Anti-Green Fluorescent Protein (GFP) Polyclonal Antibody, Unconjugated, Molecular Probes | Invitrogen | Molecular Probes Cat# A-11122, RRID:AB_221569 | 1:500 dilution |
| Antibodies | MBP (C-16) antibody | Santa Cruz | Santa Cruz Biotechnology Cat# sc-13914, RRID:AB_648798 | 1:500 dilution |
| Antibodies | Cy3-AffiniPure Goat Anti- | Jackson ImmunoResearch Labs | Jackson ImmunoResearch Labs Cat# 111- | 1:200 dilution |
|  | Rabbit IgG (H+L) antibody |  | 165-003, RRID:AB_2338000 |  |
| Antibodies | Cy3-AffiniPure Donkey Anti-Goat IgG (H+L) antibody | Jackson ImmunoResearch Labs | Jackson ImmunoResearch Labs Cat# 705-165-003, RRID:AB_2340411 | 1:200 dilution |
| Antibodies | Alexa Fluor 488-AffiniPure Donkey Anti-Rabbit IgG (H+L) antibody<br>Donkey anti-Rabbit IgG (H+L) ReadyProbes Secondary Antibody, Alexa Fluor 488 | Jackson ImmunoResearch Labs | Jackson ImmunoResearch Labs Cat# 711-545-152, RRID:AB_2313584 | 1:200 dilution |
| Software | GraphPad Prism software | GraphPad Prism (https://graphpad.com) | RRID:SCR_002798 | Version 8.0.0 |
| Software | Fiji | https://imagej.net/Fiji | RRID:SCR_002285 |  |
| Software | ANTs (Advanced Normalization ToolS) | http://stnava.github.io/ANTs/ | RRID:SCR_004757 |  |
| Software | FSL | http://www.fmrib.ox.ac.uk/fsl/ | RRID:SCR_002823 |  |
| Software | Python | http://www.python.org/ | RRID:SCR_008394 |  |
| Software | SPM | http://www.fil.ion.ucl.ac.uk/spm/ | RRID:SCR_007037 |  |
| Software | Nitime | http://nipy.org/nitime/ | RRID:SCR_002504 |  |
| Software | ITK-Snap | http://www.nitrc.org/projects/itk-snap/ | RRID:SCR_002010 |  |
| Software | TrackVis | http://www.trackvis.org/ | RRID:SCR_004817 |  |
| Software | SMART Video tracking, Panlab | https://www.panlab.com/en/products/smart-video-tracking-software-panlab | RRID:SCR_002852 |  |
| Software | Doric Neuroscience studio Software | http://doriclenses.com/life-sciences/software/955-doric-neuroscience-studio.html | Doric Neuroscience Studio, RRID:SCR_018569 |  |
| Software | Metofico Photometry Analyzer | https://metofico.com/ | Metofico Photometry Analyzer_v3 |  |

### Animals

Experiments were performed on 16∼22-week-old male R6/1 transgenic mice (B6CBA background) expressing the exon-1 of mutant huntingtin with 115 CAG repeats and their age-matched wild-type littermate as controls. Mice were acquired from The Jackson Laboratory (Bar Harbor, ME) and housed together in groups of mixed genotypes with access to food and water ad libitum in a colony room kept at 19-22°C and 40-60% humidity, under a 12:12 hr light/dark cycle. Genotypes were determined by PCR from ear biopsy, mice randomly assigned to experimental groups and data were recorded for analysis by microchip mouse number. A power analysis was performed for fMRI and the behavioural experiments based on previous data from the laboratory, considering power of 0.8, alpha of 0.05 and standard deviation of 20%, which leads to N=12 for behaviour. N values (biological replicates) are given throughout the manuscript in the figure legends. Each experiment was performed once. All animal procedures were conducted in accordance with the Spanish RD 53/2013 and European 2010/63/UE regulations for the care and use of laboratory animals and approved by the animal experimentation Ethics Committee of the Universitat de Barcelona and Generalitat de Catalunya.

### MRI Image acquisition

Experiments were blindly performed on a 7.0T BioSpec 70/30 horizontal animal scanner (Bruker BioSpin, Ettlingen, Germany), equipped with an actively covered gradient structure (400 mT/m, 12 cm inner diameter). To evaluate connectivity between regions of interest, mice (*n*=11 WT, *n*=13 R6/1) at 17-20 weeks of age underwent structural T2-weighted imaging, diffusion weighted imaging (DWI) and rs-fMRI. Animals were placed in a Plexiglas holder in a supine position and were fixed using tooth and ear bars and adhesive tape; a combination of anesthetic gases [medetomidine (bolus of 0.3 mg/kg, 0.6 mg/kg/h infusion) and isoflurane (0.5%)] was administered using a Plexiglas holder with a nose cone.

Proper position of the head at the isocenter of the magnet was ensured with a 3D-localizer scan. T2-weighted image was obtained using a RARE sequence [effective TE = 33 ms, TR = 2.3 s, RARE factor = 8, voxel size = 0.08 x 0.08 mm2, slice thickness = 0.5 mm]. DWI was acquired using an echo planar imaging (EPI) sequence [TR = 6s, TE = 27.7 ms, voxel size 0.21 x 0.21 mm^2^, slice thickness = 0.5 mm, 30 gradient directions with b = 1000 s/mm^2^ and 5 baseline images without diffusion weighting]. rs-fMRI acquisition was performed using an EPI sequence [TR = 2 s, TE = 19.4 ms, voxel size 0.21 x 0.21 mm2, slice thickness = 0.5 mm]; 420 volumes were acquired and the total scan time was 14 min. After completion of the imaging session, atipamezol (Antisedan®, Pfizer) and saline were injected to reverse the sedative effect and compensate fluid loss.

### Functional connectivity analysis

Seed-based analysis was performed as previously (Fernández-García et al., 2020) to evaluate functional connectivity of the M2 cortex with the rest of the brain. Pre-processing of the rs-fMRI data included: slice timing, spatial realignment using SPM8 for motion correction, elastic registration to the T2-weighted volume using ANTs for EPI distortion correction (Avants et al., 2008), detrend, smoothing [full-width half maximum (FWHM) = 0.6 mm], frequency filtering of time series between 0.01 and 0.1 Hz and regression by motion parameters using NiTime (http://nipy.org/nitime).

Brain parcellation was defined according to the MRI-based atlas of the mouse brain (Ma et al., 2008). The sensorimotor cortex defined in the original atlas was manually divided into somatosensory cortex (Som), orbitofrontal cortex (OFC), primary motor (M1) cortex, M2 cortex, cingulate cortex (Cg), medial prefrontal cortex (mPFC) and retrosplenial cortex (RS). The atlas template was elastically registered to each subject T2-weighted volume using ANTs (Avants et al., 2008) and finally applied to the label map to obtain specific parcellation for each animal.

Brain parcellation was registered from T2-weighted to resting state fMRI images. The left M2 cortex was selected as seed region for seed-based analysis. Extracted average time series in the M2 cortex was correlated with the time series of all the voxels within the brain, resulting in a connectivity map containing the value of the correlation between the BOLD (blood-oxygen level dependent) signal time series in each voxel with the seed time-series. Seed to region connectivity was calculated as the mean value of the correlation map in each region, considering only positive correlations.

### Structural connectivity analysis

The fiber tracts connecting M2 cortex with SC were identified and characterized based on automatically identified regions in the T2-weighted image and tractography results from DWI. Diffusion tensor images (DTI) were estimated and standard DTI metrics were computed, including Fractional Anisotropy (FA), Mean Diffusivity (MD), Axial Diffusivity (AD) and Radial Diffusivity (RD). To estimate the fiber tracts, DWI was preprocessed, including eddy-current correction using FSL(Jenkinson et al., 2012), denoising (Coupe et al., 2008) and bias correction(Tustison et al., 2010). The five baseline images were averaged and registered to the T2-weighted image to correct for EPI distortions. Whole brain tractography was performed using a deterministic algorithm based on constrained spherical deconvolution model, considering as seed points the voxels where FA>0.1. The same threshold was defined as stop criterion for the algorithm. Once the fiber tracts were identified, its average FA, MD, RD and AD were computed.

### Stereotaxic surgery

We used different adeno-associated virus (AAV): AAV-YFP virus construct under CamKII promotor (AAV1-CaMKIIa-eYFP-WPRE.hGH) was used for viral tracing characterization; AAV-ChR2 virus construct under CamKII promotor (AAV1-CaMKIIa-hChR2(H134H)-eYFP-WPRE.hGH) was used for optogenetic modulation; AAV-GCaMP6f virus construct under SYN promotor (AAV9-Syn-GCaMP6f-WPRE-SV40) was used for fiber photometry analysis. Virus production, amplification, and purification were performed by University of Pennsylvania-Penn Vector Core and Addgene (titers: ∼1 x 10^12^ genomic particles/mL).

Stereotaxic surgery was performed under isofluorane anesthesia (5% induction and 1.5% maintenance) in 16-week-old mice. A volume of 0.5µl of corresponding viral constructs was injected in M2 cortex [mm from bregma and dura matter: +2.46 anteroposterior (AP), ±1 mediolateral (L), -0.8 dorsoventral (DV)] using 5 µl Hamilton syringe with a 33 gauge needle at 0.1 µl/min. To allow diffusion of virus particles and avoid refluxes, needle was left for an additional 5 min period. Animals were housed for viral expression and recovery from surgery at least 4 weeks before experiments were initiated.

For optogenetic stimulation in vivo, fiber-optic cannulas (MFC_200/240-0.22_2.5_ZF1.25_FLT; Doric Lenses) were implanted bilaterally in the superior colliculus [mm from bregma and skull: AP -4.04, L ±1.45, DV -1.75]. For fiber photometry analysis in vivo, fiber-optic cannulas (MFC_400/430-0.66_1.0mm_MF1.25_FLT; Doric Lenses) were implanted unilaterally in the left hemisphere of the M2 cortex [mm from bregma and dura matter: +2.46 anteroposterior (AP), ±1 mediolateral (ML), -0.8 dorsoventral (DV)]. All fiber-optic cannulas were implanted during surgery and fixed in place using dental cement.

### Behavioural assessment

All behavioural tests were performed by an experimented observer during the light phase and animals were habituated to the experimental room for at least 1 hour before testing. All testing apparatus were cleaned with water and dried between tests and animals.

#### Beetle Mania Task (BMT)

The beetle mania task (BMT) assesses both passive and, in particular, active fear responses of the mice to a unexpected moving beetle (Nano Nitro, Hexbug) (Almada et al., 2018; Heinz et al., 2017). The test was performed using a white rectangular arena (15 x 40 x 30 cm) with dim light (∼20 lux). Briefly, the test comprises two successive phases of 5 min: during the habituation phase, mice freely explored the arena. During this time, distance travelled and rearing times were scored using the SMART 3.0 software (Panlab). In the testing phase, confrontations with the erratically moving robo-beetle were scored. The robo-beetle was introduced to the arena at maximal distance to the mouse and the following behavioural responses analysed: (1) escape, mice withdrawals with accelerated speed in the direction opposite to the beetle’s movement vector; (2) tolerance, ignorance of the robo-beetle after physical contact; (3) approach, mice follow the robo-beetle in close vicinity; (4) avoidance, mice moves in opposite direction to the robo-beetle and (5) total contacts, the sum of all the events evaluated. Escape events were normalized to total contacts, whereas tolerance, approach and avoidance were normalized to the sum of these three parameters, without the escape events. For the experiment including optogenetic stimulation, the testing phase was prolonged an additional 5 min when stimulation was performed.

#### Looming task

The looming task assesses light-triggered behaviours in freely moving mice and was performed based in (Liang et al., 2015). The arena consisted in a narrow and long hall (8 x 150 x 30 cm^3^) dimly illuminated (∼20 lux) with a bulb with bright and white light placed at equity distance from the beginning and the end of the corridor. Mice were placed in one site of the hall and freely walked to the opposite site, then, when passed below the bulb it triggered a ∼1 s light flash above the animal. Each animal was tested for 5 trials, with inter-trial intervals of ∼20 min, and scored: time 1, time mice walked from the beginning of the hall to the bulb; and time 2, time mice walked from the bulb to the end of the corridor.

### Optogenetic stimulation

473 nm diode-pumped solid-state blue laser (Laserglow) was used to deliver blue light [10 Hz, 5 ms pulse width, ∼5 mW (at tip of each fibre)] using a custom-made wave-form generator (Arduino). For electrophysiological recordings ex vivo, fiber-optic patch cord was placed above the slice surface. For in vivo experiments, light beam was split using a rotary joint (FRJ 1×2) and patch cords (200 µm core, 0.22 NA) were connected to the implanted fiber-optic cannulas with Zicronia sleeves (Doric Lenses). Mice were habituated to manipulation before the experiment.

### Multi-electrode arrays (MEA)

Electrophysiological recordings were performed as previously described (Fernández-García et al., 2020; Kim et al., 2020). Briefly, coronal sections of mouse brain were obtained on a vibratome at 350 µm thickness in ice-cold artificial cerebrospinal fluid (aCSF) oxygenated (95% O_2_, 5% CO_2_) and then transferred for 15 min to an oxygenated 32°C recovery solution. Slices were then transferred to oxygenated aCSF at room temperature for at least 1 hr before recording.

Electrophysiological data were recorded by a blind experimenter using a MEA set-up from Multi Channel Systems MCS GmbH (Reutlingen, Germany). MC Stimulus and MC Rack software from Multi Channel Systems were used for stimulation, recording, and signal processing. We assessed the correct position of the slices on the electrode field using a digital camera during the recording.

Field post-synaptic currents (fPSC) were recorded in the lateral SC in response to the stimulation of M2 cortical afferents with 1 ms 473 nm light pulses of increasing intensities (0.024, 0.033, 0.911, 4.23, 10.2, 19.8, 44.8 and 79.0 mW/mm^2^). The laser was approximately placed in -4.04 posterior to bregma, -1.75 ventral from the skull, and ±1.45 lateral from the middle, corresponding to lateral SC. Evoked fPSCs responses in the SC were analyzed after trains of 1 ms light stimuli.

For electrical stimulation, we set one MEA electrode located lateral to the SC and generated input/output curves by trains of three positive-negative identical pulses at increasing currents (250 – 3000 µV).

### Fiber photometry

Changes of neuronal activity were assessed using the GCamp6f fluorescent calcium indicator and normalized using isosbestic fluorescent recordings (405nm), using a custom made fiber photometry system obtained from Doric Lenses. We used the free Doric Neuroscience studio Software to control the Doric console and LED drivers. A 465-nm LED (CLED_465, Doric Lenses) modulated at 241.58 MHz and a 405_nm LED (CLED_405, Doric Lenses) modulated at 572.21 MHz were directed and coupled into a personalized fluorescence mini cube (iFMC7, Doric Lenses) through 1 m attenuator fiberoptic patchcords (400 µm core, 0.48 NA). Combined 465_nm and 405_nm light were launched to a pigtailed (400 µm core, 0.57 NA) fiberoptic rotary joint (Doric Lenses) connected to a low autofluorescence mono fiber-optic patchcord (400 µm core, 0.57 NA) allowing freely moving tests with mice, and finally mated to the implanted fiberoptic cannula (see stereotaxic surgery) via a mating black covered Zicronia sleeve. Emitted fluorescence travels back through the mini cube and spectra is detected and amplified by the Fluorescence Detector Amplifier from Doric and data is send to the fiber photometry console. Bleaching of the fiberoptic patchcords will be done prior to each experiment for 3 h. Mice were connected to the optical fiber 5 min before the task and the 635 and 405 LED were turned on to ensure stable photometric recordings.

Raw and demodulated 465 nm and 405 nm fluorescent intensity data is recorded at 12000 Hz in freely-moving mice using the free Doric Neuroscience studio Software. Demodulated data is further processed by a custom made software provided by Metofico and based on Matlab. Briefly, data is down sampled to 1 Hz and artifacts are removed using a custom digital filter. Average fluorescence during one minute window before the introduction of the beetle was considered as baseline (ΔF/F_0_). To correct movement artifacts, normalized 405 nm signal was subtracted from the normalized 465 nm.

### Immunohistochemistry

Mice were sacrificed by cervical dislocation and brains were post-fixed with 4% paraformaldehyde for immunohistochemical analysis, dehydrated in a PBS/Sucrose gradient [from 15% (48 hr post-mortem) to 30% (32 hr post-mortem)] with 0.02% Sodium Azide, snap frozen and stored at −20°C. Sagittal sections (25 µm sagittal) were cut on a cryostat (Microm) and preserved in PBS with 0.02% Sodium Azide at 4°C. Anti-GFP (1:500, Invitrogen, #11122) antibody was used to evaluate expression of AAV-YFP, AAV-GCAMP6f and the AAV-ChR2 virus constructs. To quantify M2 cortical afferents to the SC, anti-MBP (Myelin Binding Protein; 1:500, Santa Cruz, #sc-13914) antibody was added as axonal control. Free-floating sections were washed in PBS and permeabilized and blocked for 15 min in PBS containing 0.3% Triton X-100 and 3% normal goat serum (Pierce Biotechnology). Sections were washed again in PBS and incubated overnight at 4°C with primary antibodies. Brain slices were washed, incubated for 2 hr with the secondary antibody (1:200 goat anti-rabbit Cy3 for only GFP visualization or 1:200 donkey anti-rabbit Alexa Fluor 488 and donkey anti goat Cy3 for GFP and MBP immunofluorescence, and mounted on microscope slides using DAPI Fluoromount-G (SouthernBiotec). Fluorescence images were acquired by an epifluorescence microscope (DMI6000 Leica). No signal was detected in absence of primary antibody. Fluorescence intensity was measured using Fiji in regions of interest (ROIs). Mice with no GFP expression observed, indicating no expression of AAVs, were discarded.

### Statistical analysis

All the results were expressed as mean ± SEM. Data from individual mouse is represented by single points when possible. Statistical analysis was performed using the Student’s t-test and the two-way ANOVA analysis, followed by Bonferroni post-hoc test when appropriate, and indicated in the results section and/or figure legends. Values of *p*<0.05 were considered as statistically significant.

## Acknowledgements

We are very grateful to Ana López, Maria Teresa Muñoz and Albert Coll for their excellent technical support. We are also grateful to the staff of the Confocal Microscopy Service of the Scientific and Technological Centers of the University of Barcelona (CCiTUB). We are indebted to the Magnetic Resonance Imaging Core Facility of the Institut d’Investigacions Biomèdiques August Pi I Sunyer for the scientific technical support in MRI acquisition and analysis.

## Data and materials availability

All data needed to evaluate the conclusions in the paper are present in the paper and/or the Supplementary Materials.

